## Supplementary Information for "MIMIC: a flexible pipeline to register and summarize IMC-MSI experiments"

December 9, 2025

### 1 Supplementary

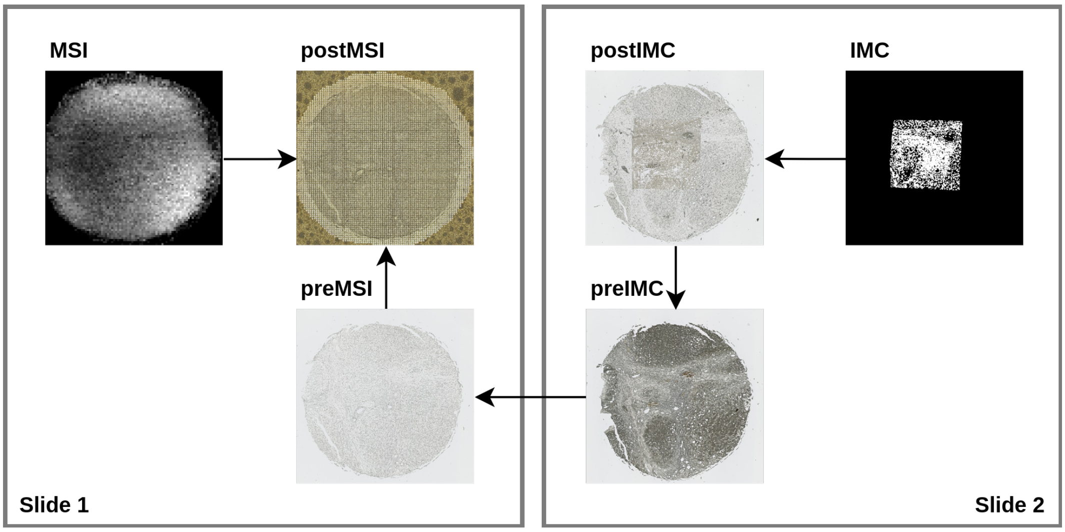


Figure S1: Broad overview of registration workflow with example images.

| **method** | **supported modalities** | **co-registration** | **co-registration validation** | **statistical methods** |
|---|---|---|---|---|
| `regToolboxMSRC` | MALDI-MSI plus histopathology | MSI to histopathology (automatic, indirect via ablation marks), histopathology to autofluorescence (automatic) | visual, landmark errors | correlation between region overlap and intensity |
| `SpaceM` | MALDI-MSI plus light microscopy | MSI to light microscopy (automatic, indirect via ablation marks) | visual, landmark errors | single-cell intensity profiles |
| `scSpaMet` | TOF-SIMS plus IMC | TOF-SIMS to IMC (automatic, direct) | visual, global intensity based | single-cell intensity profiles |
| Nunes *e*t al. | MALDI-MSI plus IMC | MALDI-MSI to IMC (manual, direct) | visual | single-cell intensity profiles |
| `msiFlow` | MALDI-MSI plus IF | MALDI-MSI to IF (automatic, direct) | visual, region overlaps | single-cell intensity profiles |
| `MIMIC` | MALDI-MSI plus IMC | MALDI-MSI to IMC (automatic, indirect via ablation marks and intermediate microscopy images) | visual, landmark errors, region overlaps | MSI pixel-level celltype compositions |

Table S1: Broad overview of computational tools for the co-registration and analysis of multi-modal MSI data.

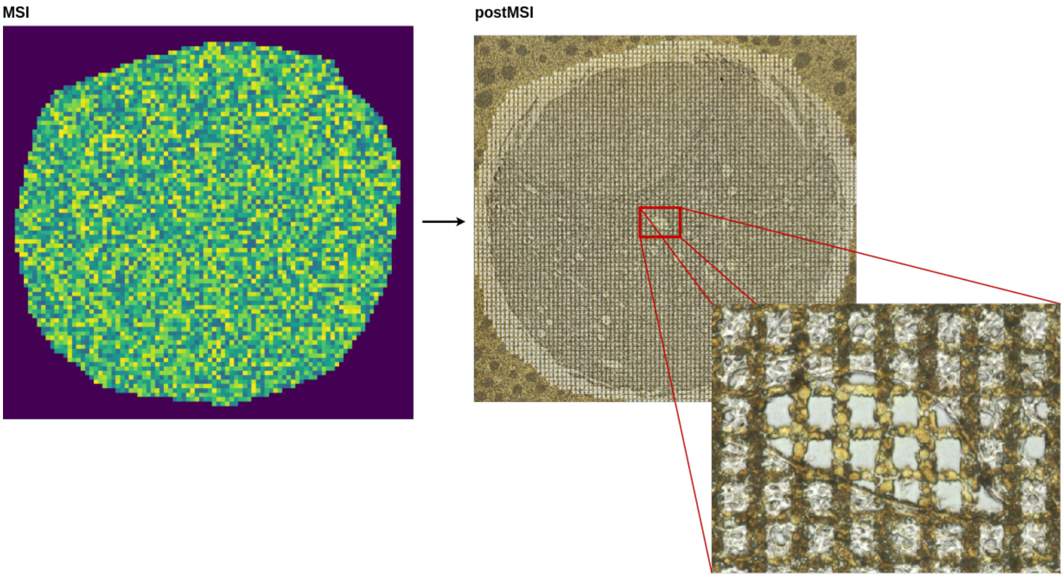


Figure S2: Example MSI to postMSI registration.

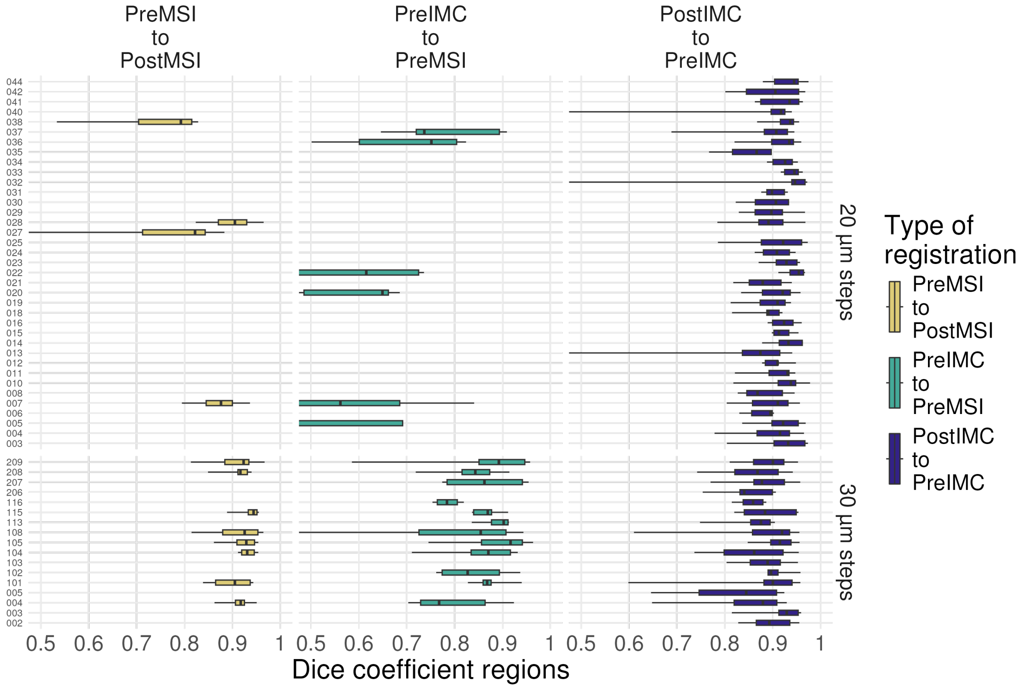


Figure S3: Evaluation precision of registrations. DICE coefficients of regions extracted from the four microscopy images and compared agains each other.

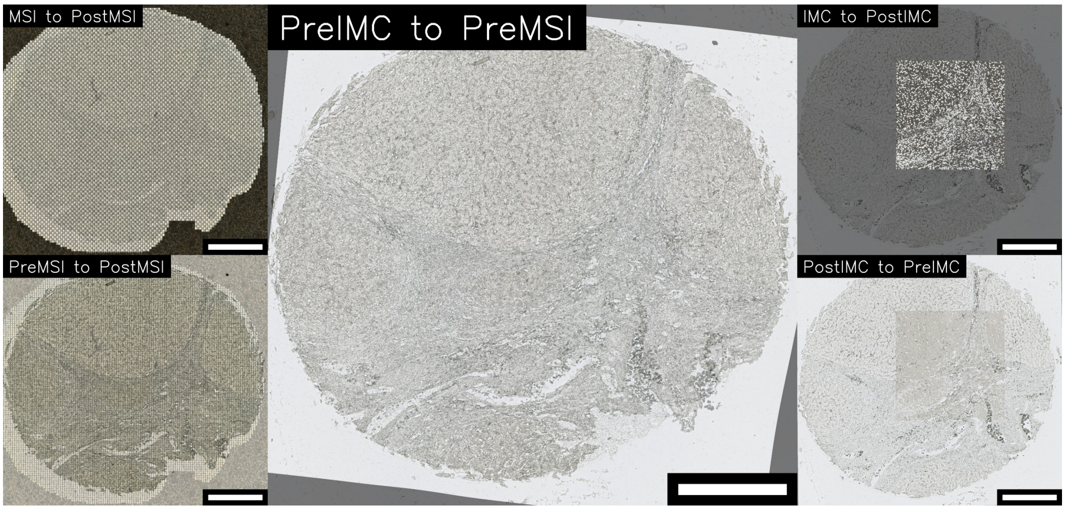


Figure S4: Blended images after co-registration for sample NASH 005.

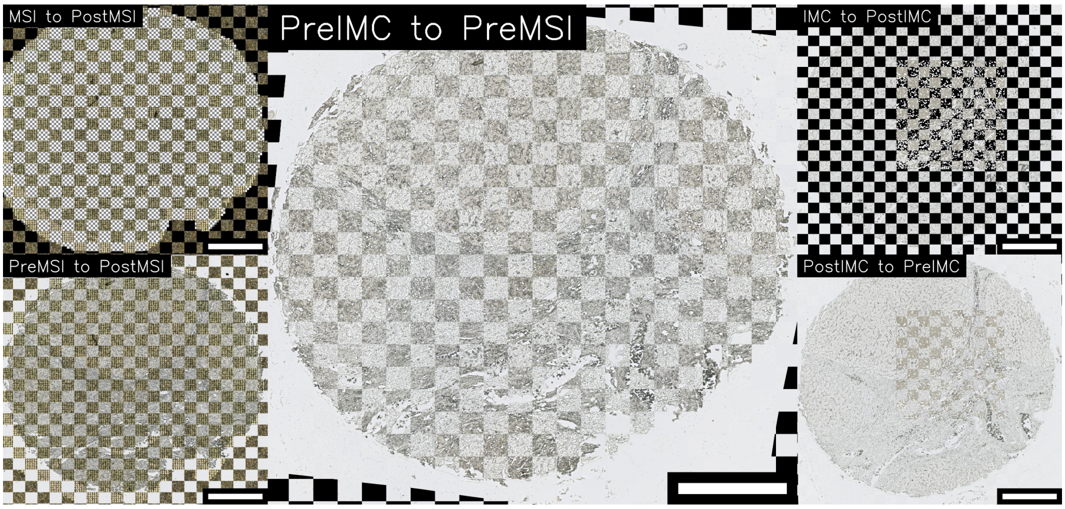


Figure S5: Checkerboard images after co-registration for sample NASH 005.

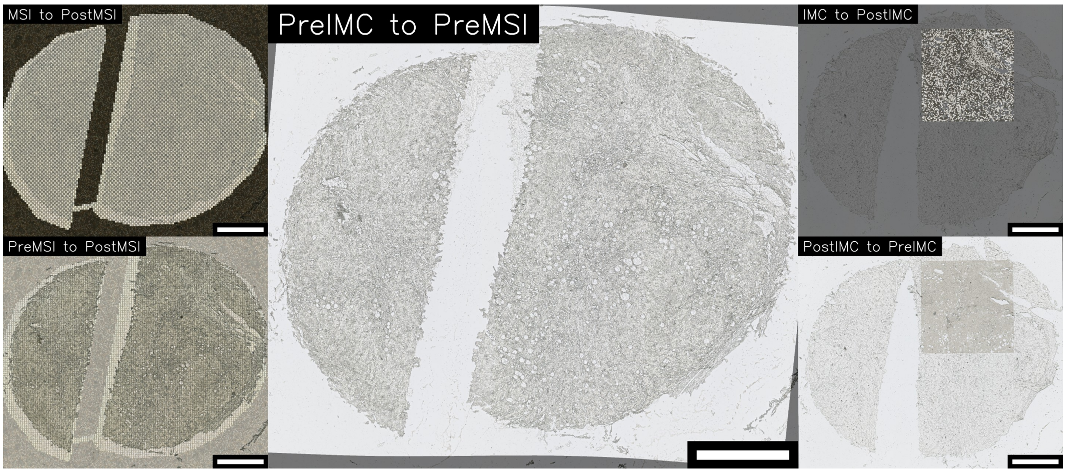


Figure S6: Blended images after co-registration for sample NASH 011.

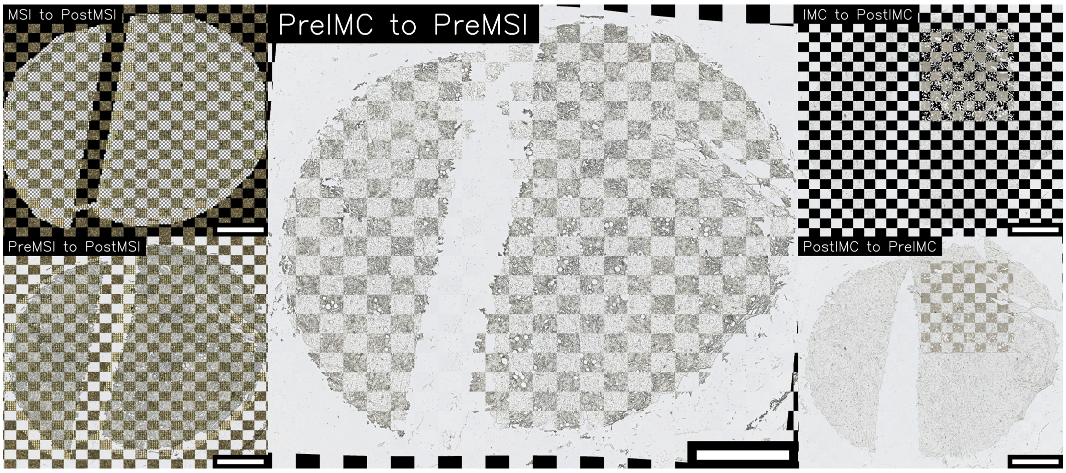


Figure S7: Checkerboard images after co-registration for sample NASH 011.

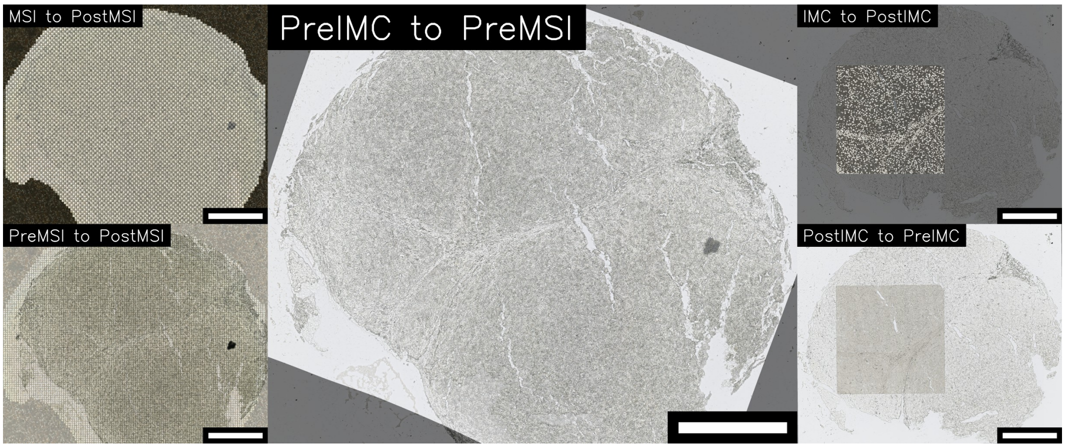


Figure S8: Blended images after co-registration for sample NASH 020.

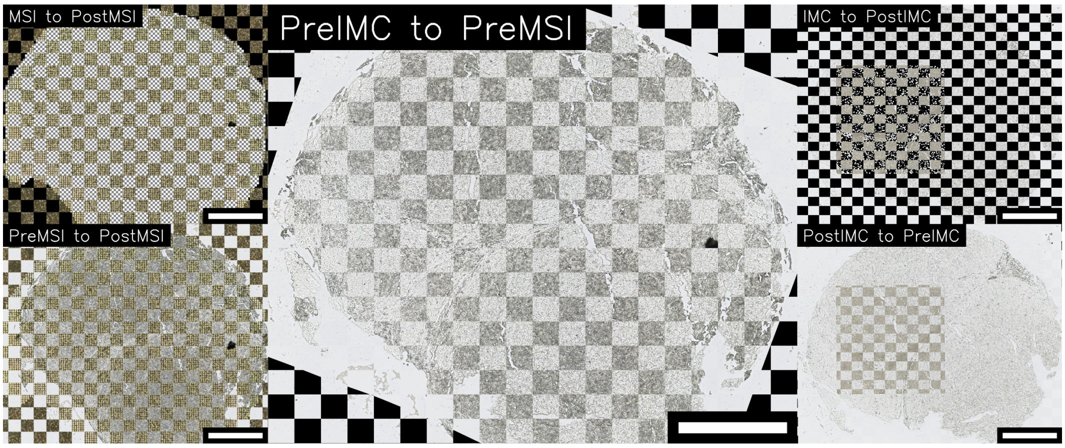


Figure S9: Checkerboard images after co-registration for sample NASH 020.

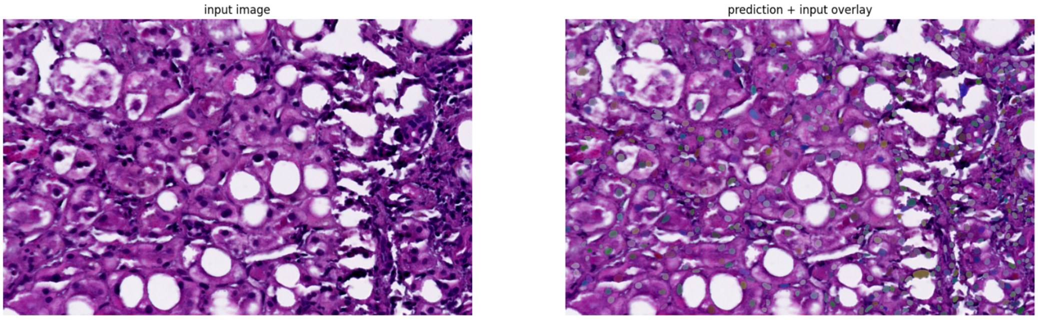


Figure S10: Example image of nuclei detection. Left image: H&E stained image, Right image: overlaid nuclei as detected with StarDist.

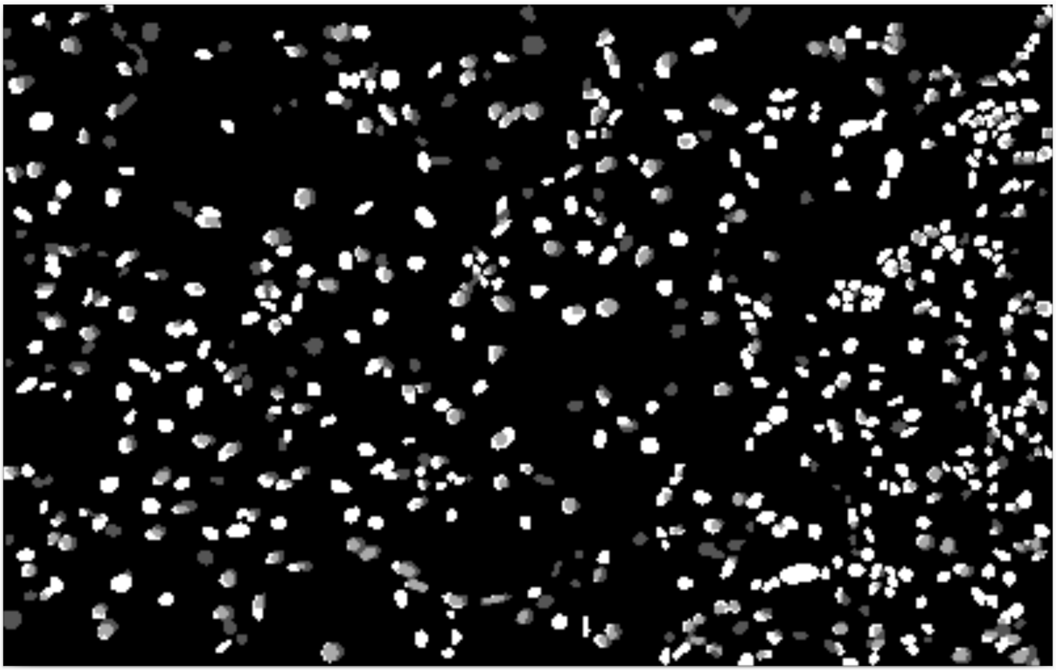

Figure S11: Example image of nuclei detection. White: nuclei from H&E image, Dark gray: nuclei from IMC, Light gray: overlap of the two masks, Black: no nuclei for either modality.

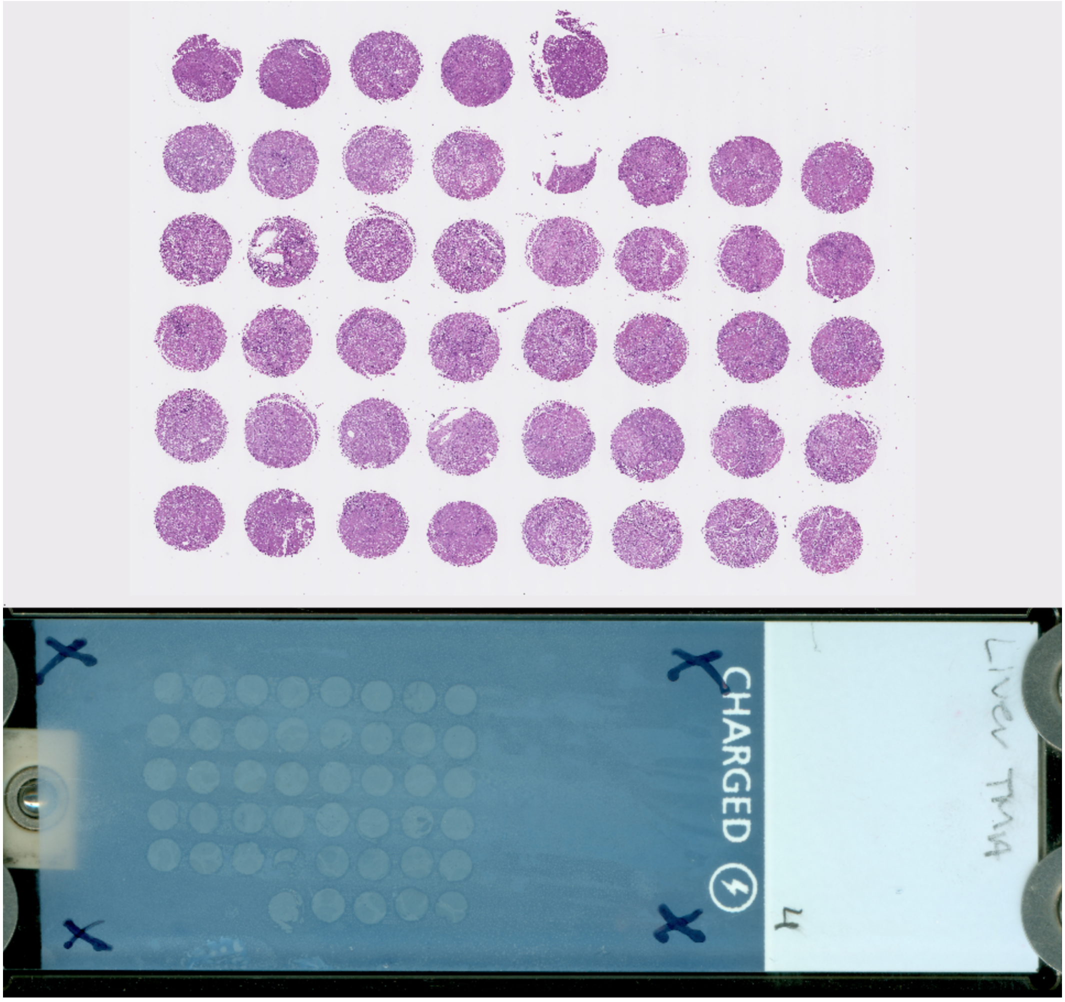


Figure S12: Images used for manual vs automatic image registration precision. Top: high resolution H&E stain, bottom: low resolution slide scan.

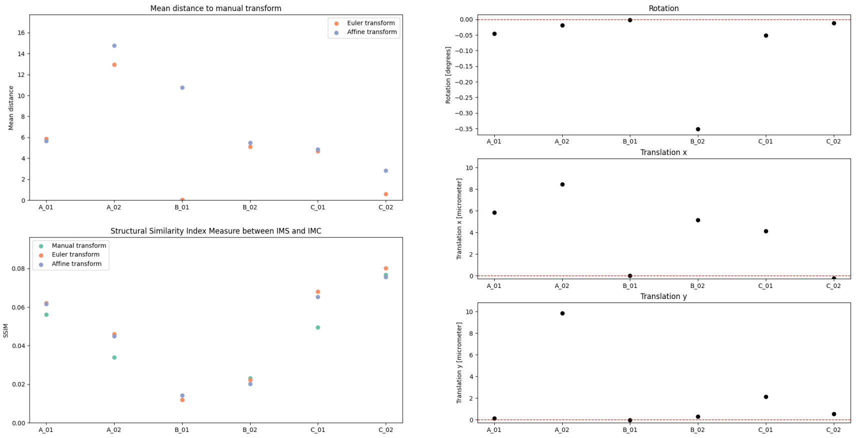


Figure S13: Registration parameters and metrics of MSI to MSI coregistration for the Nunes et al. data. Top left: Mean distance of grid point between the manual and the euler or affine transformation. Bottom left: Structural similarity index measure between IMC and MSI for the different transformations. Right: Euler transformation parameters from top to bottom: rotation, x-translation, y-translation.

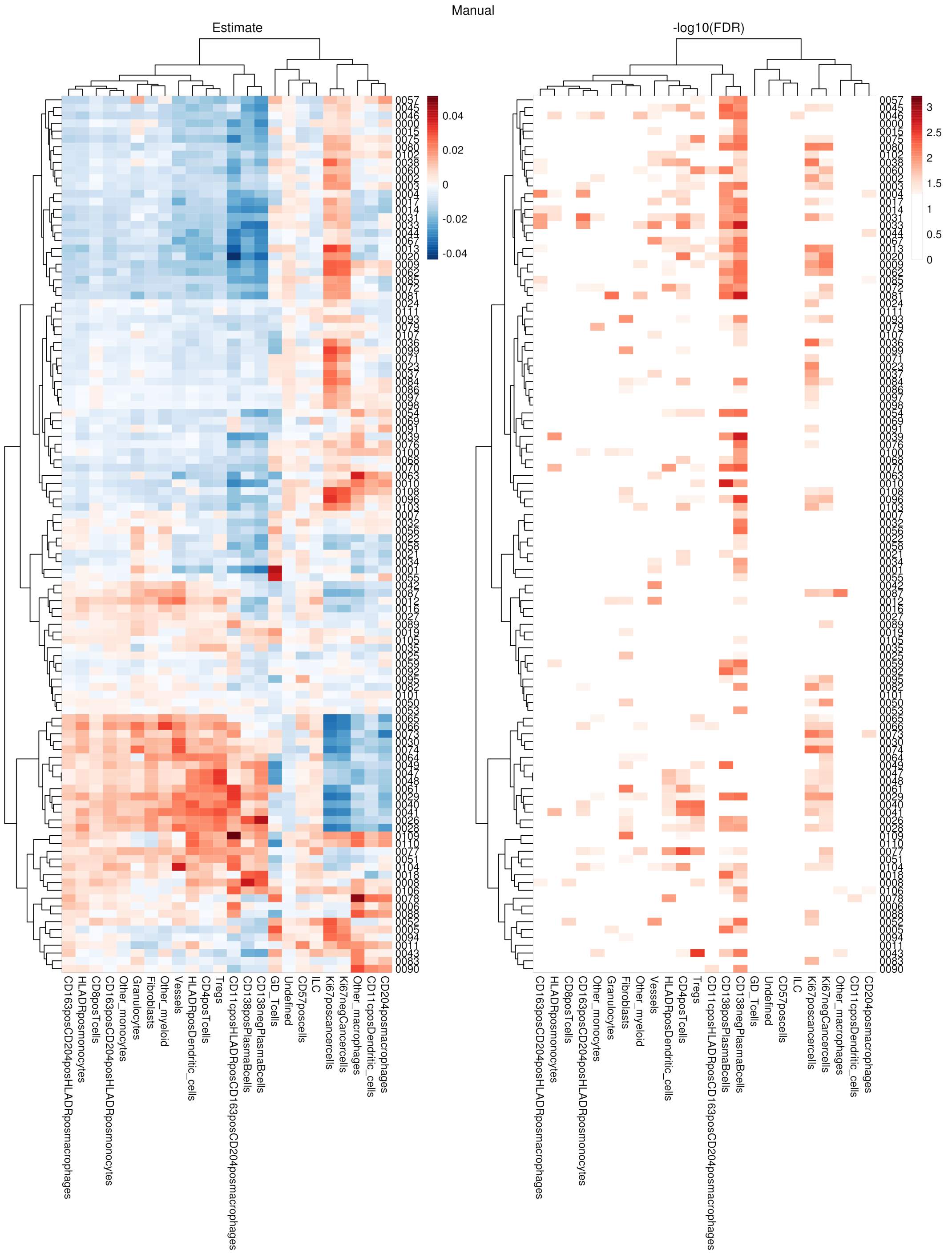


Figure S14: Association results of MSI vs. cell type proportion for the Nunes et al. data. Results for the manual transformation are shown. Left: Heatmap of estimates. Right: Heatmap of -log10(FDR corrected p-values).

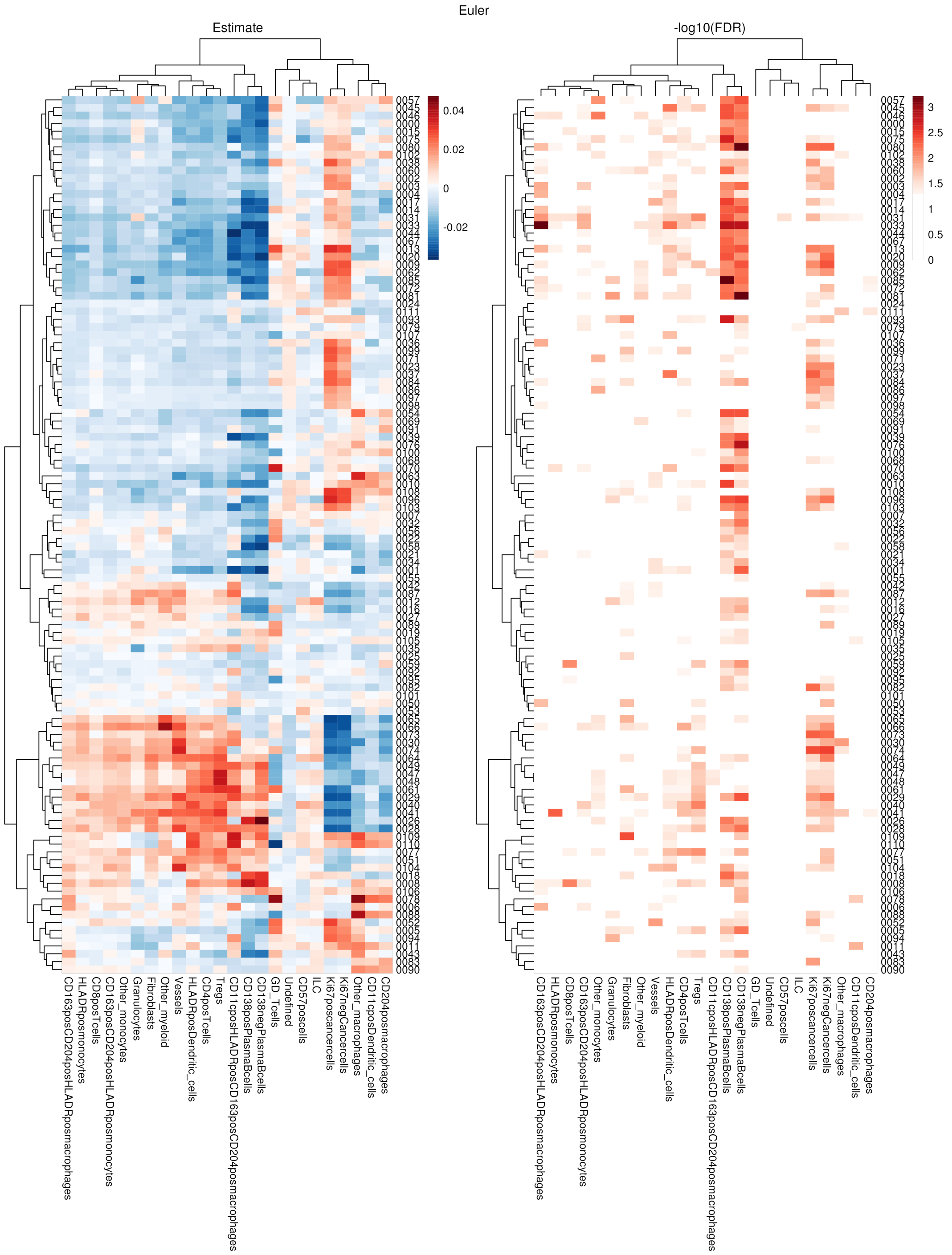


Figure S15: Association results of MSI vs. cell type proportion for the Nunes et al. data. Results for the automatic euler transformation are shown. Left: Heatmap of estimates. Right: Heatmap of -log10(FDR corrected p-values).

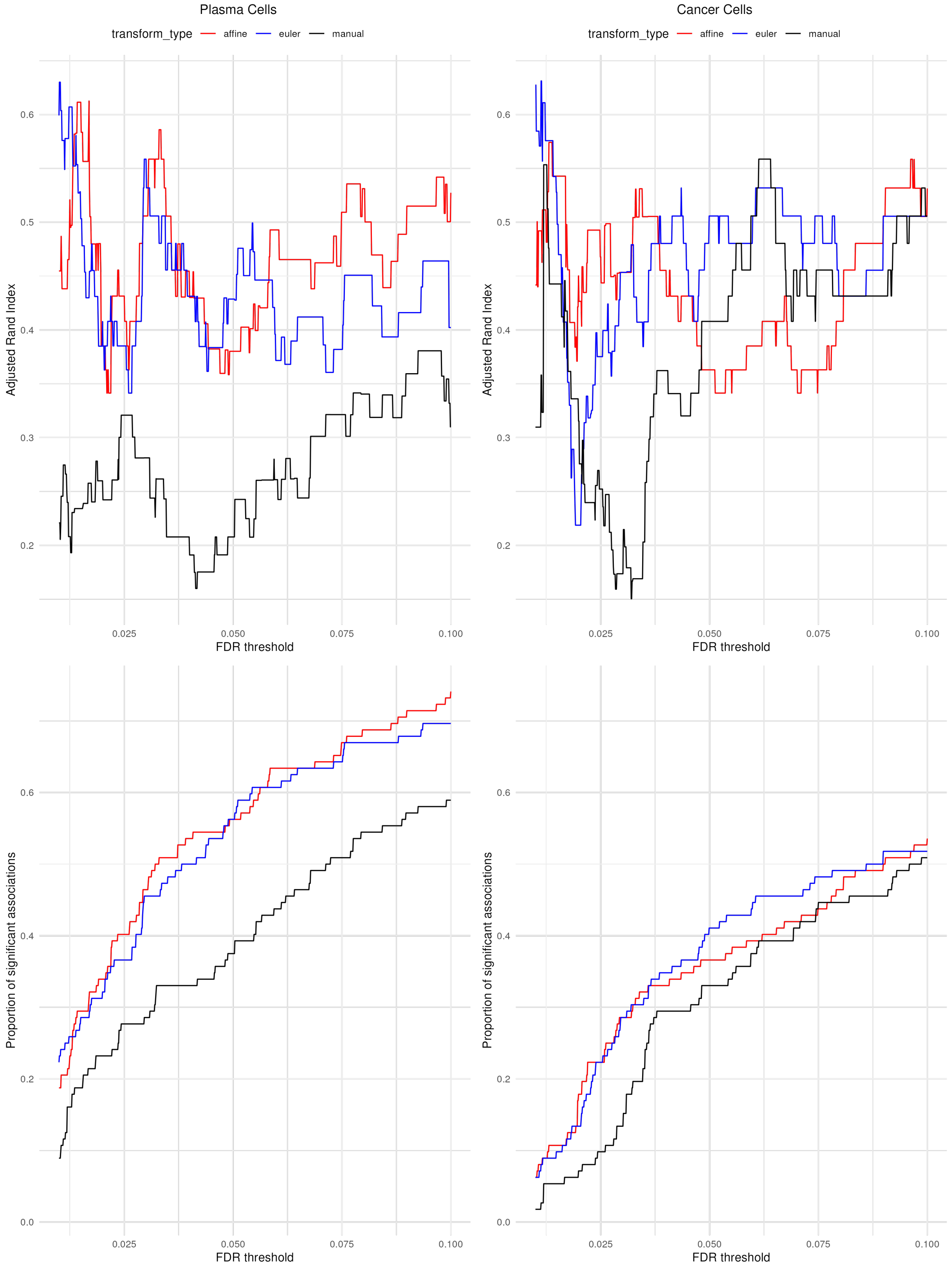


Figure S16: Comparison of modeling results of two celltypes (Plasma Cells and Cancer Cells) that both have two subcelltypes. Top row: Adjusted rand index as a function of FDR threshold, i.e. clustering m/z values into two clusters - significant and not significant - and calculation of the ARI between those two groups. Bottom row: Proportion of significant associations (i.e. both subcelltypes are significant) as a function of FDR threshold. Left column: Plasma Cells. Right column: Cancer Cells.

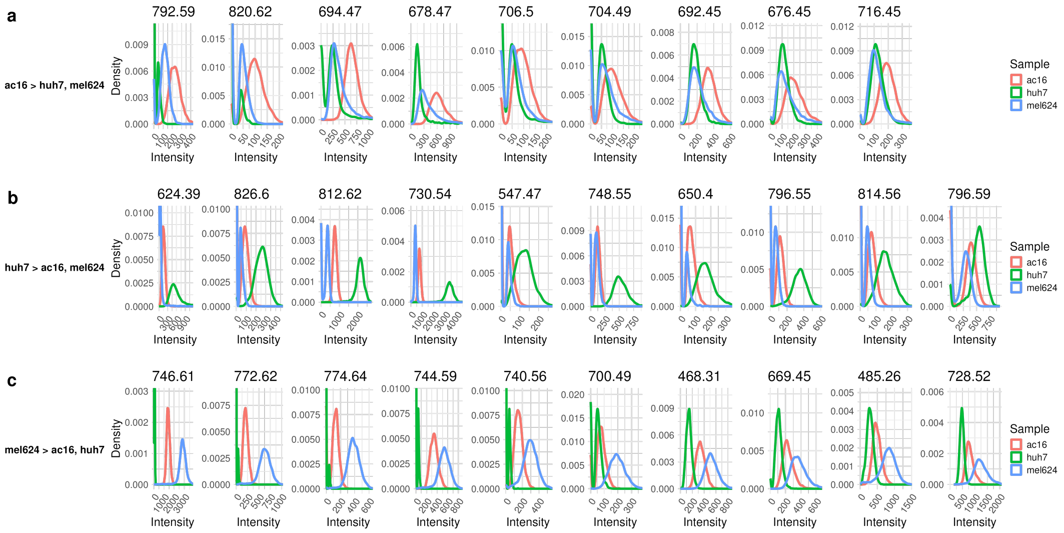


Figure S17: Distribution of peaks with different intensity between in-isolation samples. a) m/z peaks with higher intensity in cell line AC16 compared to the other two cell lines. b) higher in Huh7. c) higher in Mel624. For Huh7 only the top 10 peaks are shown.

| **Registration Step** | **Source Image** | **Target Image** | **Type of Comparison** | **Metric** | **Source Feature** | **Target Feature** |
|---|---|---|---|---|---|---|
| MSI to postMSI | MSI | postMSI | landmark | MLD | pixel coordinate | ablation mark coordinate |
| IMC to postIMC | IMC | postIMC | landmark | MLD | grid of coordinate points after transform | grid of coordinate points before transform |
| postIMC to preIMC | postIMC | preIMC | landmark | MLD | coordinate of RoMa feature | coordinate of RoMa feature |
| postIMC to preIMC | postIMC | preIMC | region | DICE | region mask from SAM | region mask from SAM |
| preIMC to preMSI | preIMC | preMSI | landmark | MLD | coordinate of RoMa feature | coordinate of RoMa feature |
| preIMC to preMSI | preIMC | preMSI | region | DICE | region mask from SAM | region mask from SAM |
| preMSI to postMSI | preMSI | postMSI | landmark | MLD | coordinate of RoMa feature | coordinate of RoMa feature |
| preMSI to postMSI | preMSI | postMSI | region | DICE | region mask from SAM | region mask from SAM |

Table S2: Registration steps and their corresponding precision evaluations. MLD: Median landmark distance.

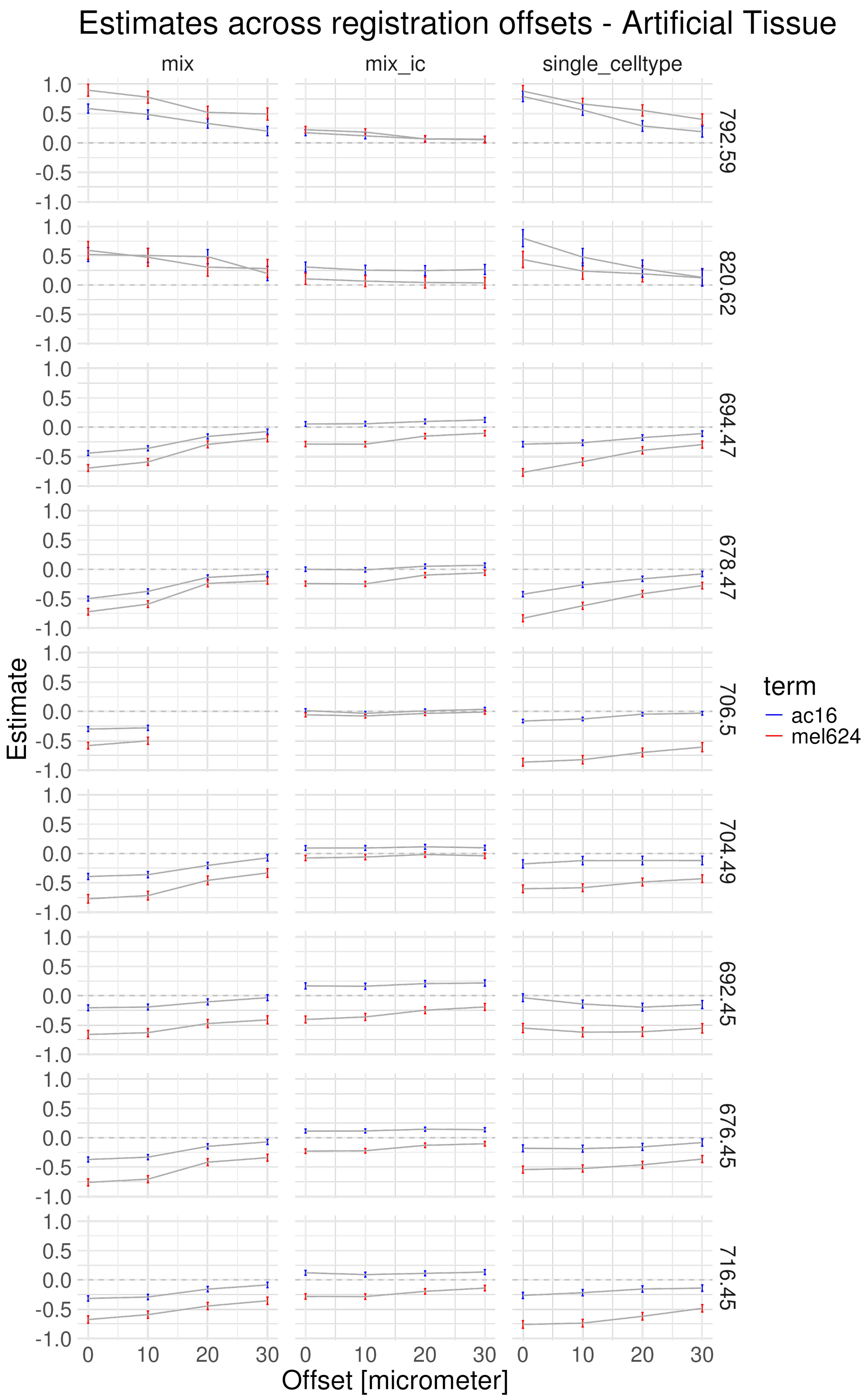


Figure S18: Sensitivity analysis of artificial tissue of m/z peaks expected to be higher in AC16 than Mel624. Offset is the number of MSI pixels that are shifted in x-axis relative to the optimal registration. mix and mix_ic are the mixed samples and single_celltype are the homogeneous samples. Shown are estimates and error bars ($+-1.96\times$ standard error).

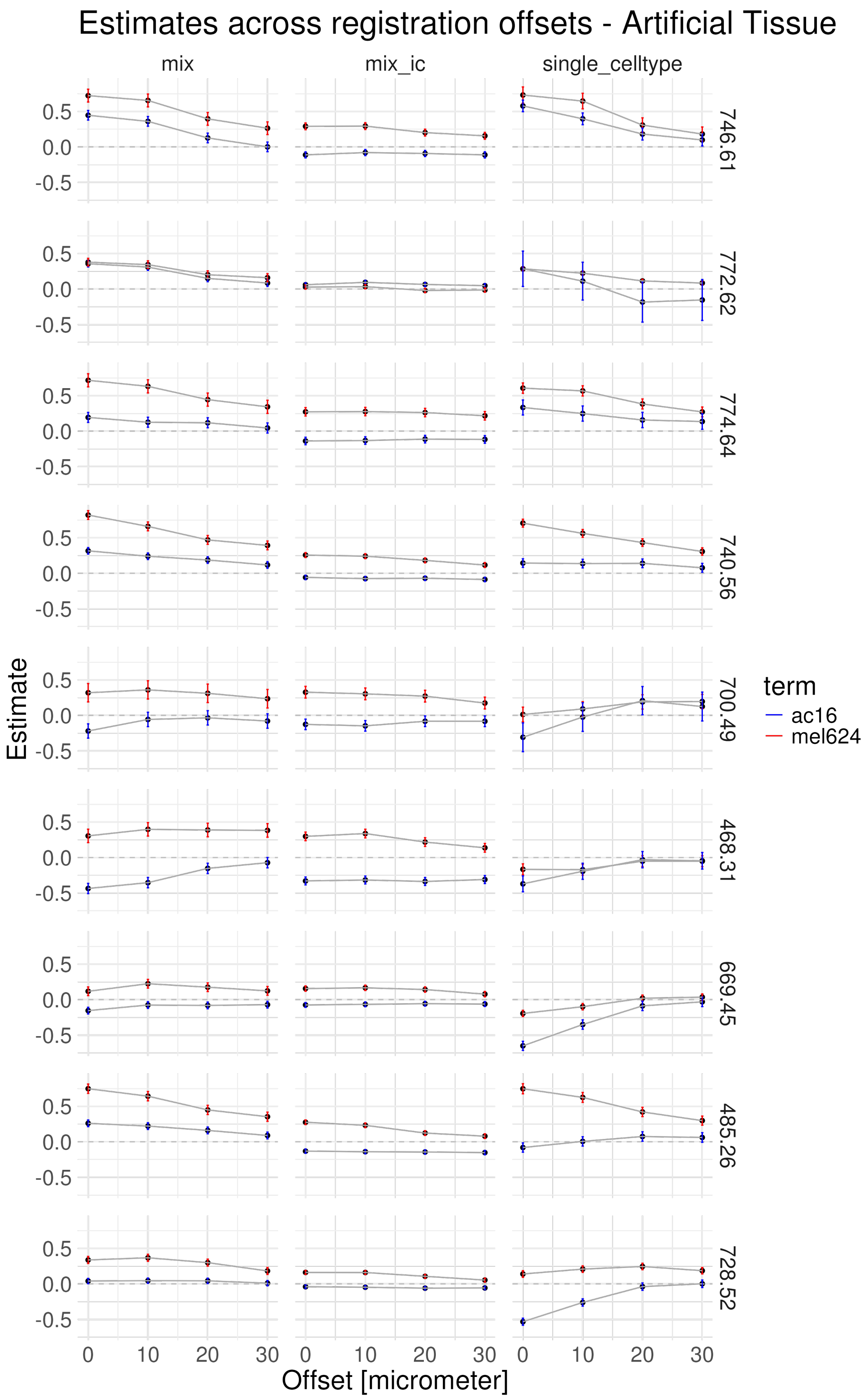


Figure S19: Sensitivity analysis of artificial tissue of m/z peaks expected to be higher in Mel624 than AC16. Offset is the number of MSI pixels that are shifted in x-axis relative to the optimal registration. mix and mix_ic are the mixed samples and single_celltype are the homogeneous samples. Shown are estimates and error bars ($+-1.96\times$ standard error).

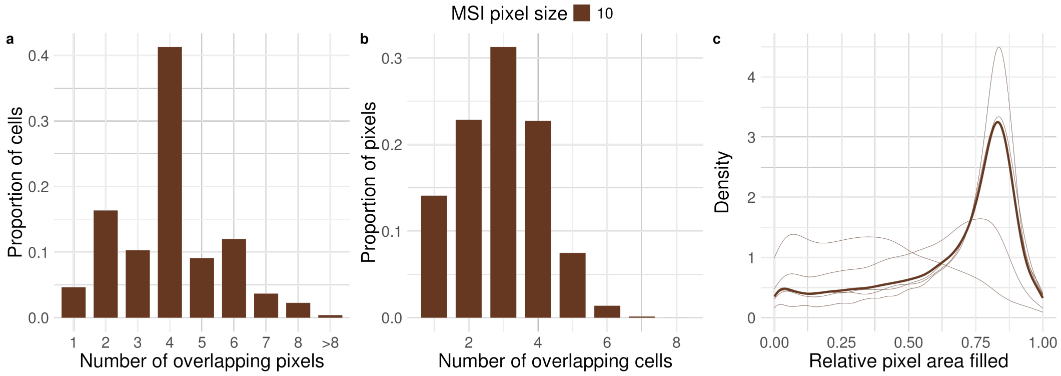


Figure S20: Distributions of overlaps between cells and MSI pixels for the artificial tissue. a) Distribution of number of pixels a cell overlaps with. b) Distribution of number of cells a MSI pixel overlaps with. c) Distribution of relative area filled by cells per MSI pixel.

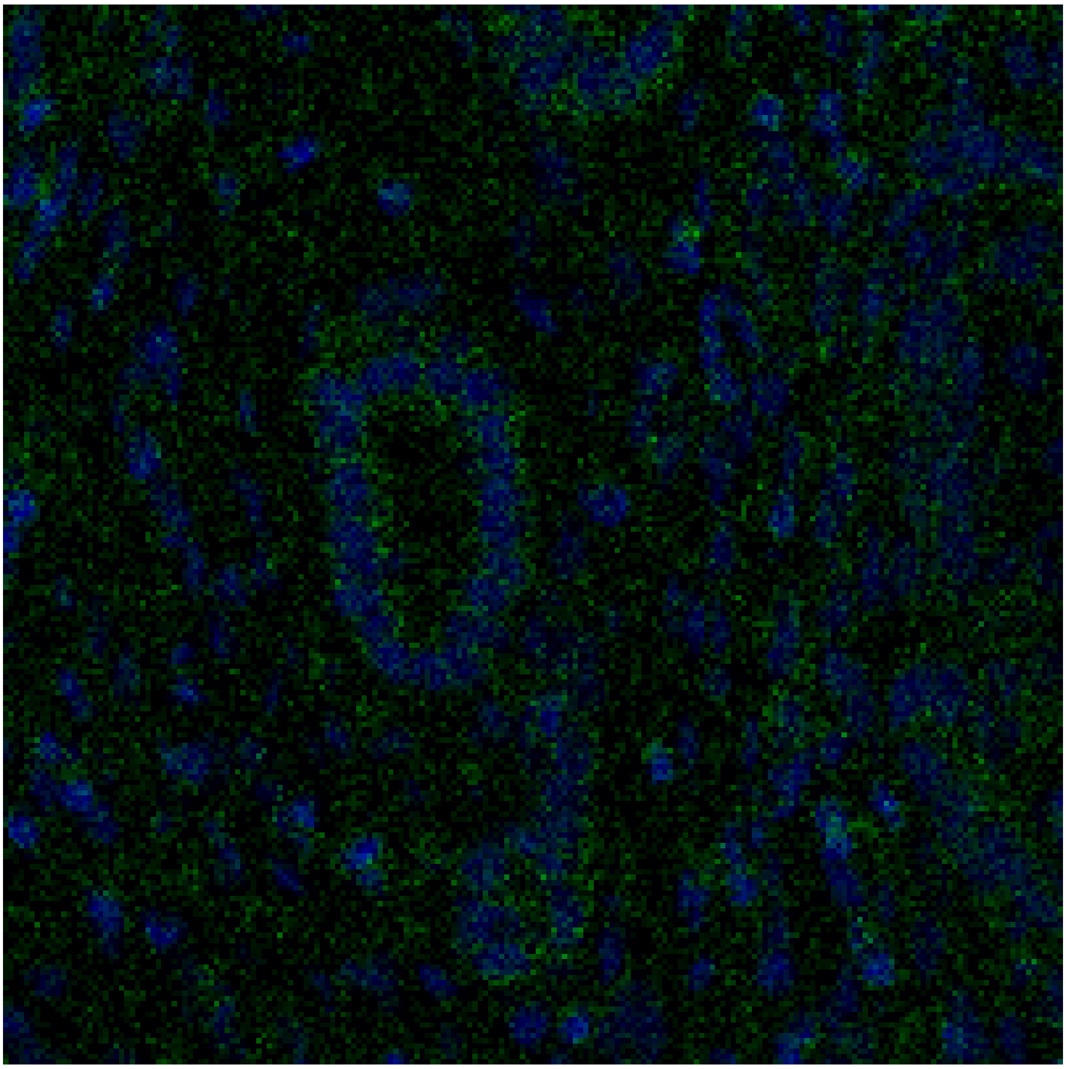

Figure S21: Example IMC image for the Liver TMA. Blue: DNA channel, green: segmentation channel.

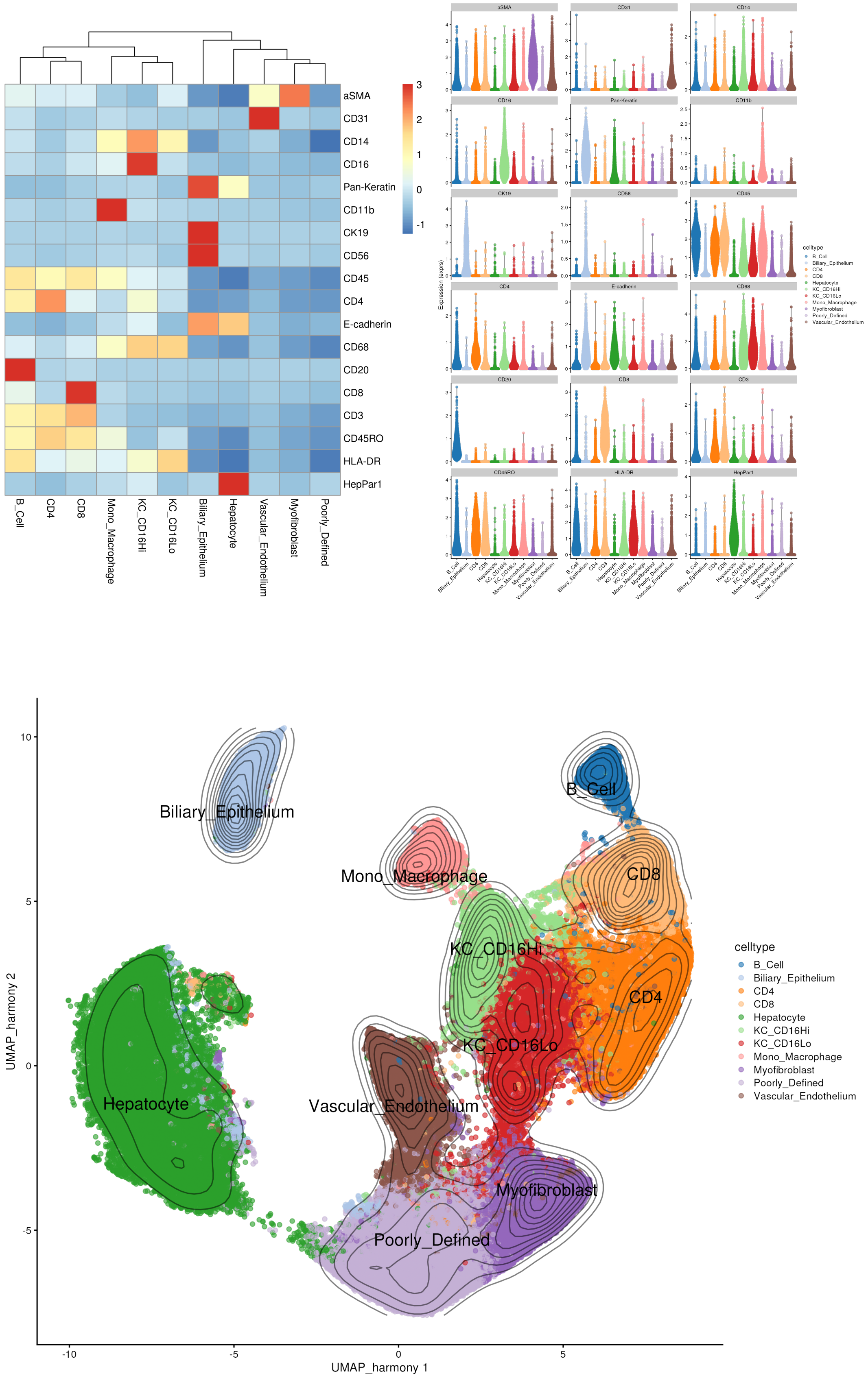




Figure S22: IMC data of Liver tissue. a) Heatmap of row-scaled mean expression per cell type. b) Violin plot of distribution of markers per cell type. c) UMAP of integrated data colored by cell type.

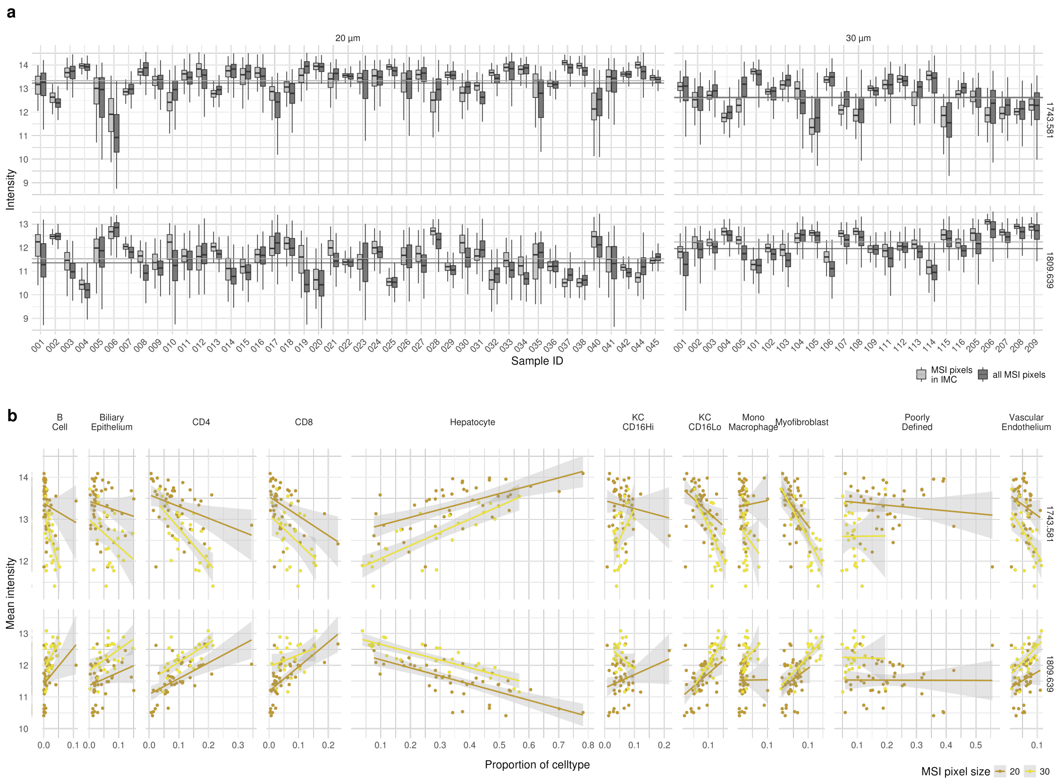


Figure S23: a) Distribution of two example m/z values (rows) for all samples (columns) colored by if all MSI pixels are used (dark gray) or if only MSI pixels on the IMC location are used (light gray). Panel titles are the MSI step size (20 or 30 microns). b) For the same two example m/z values the mean intensity vs. the cell type proportion. Line are the linear regression fit per MSI step size.

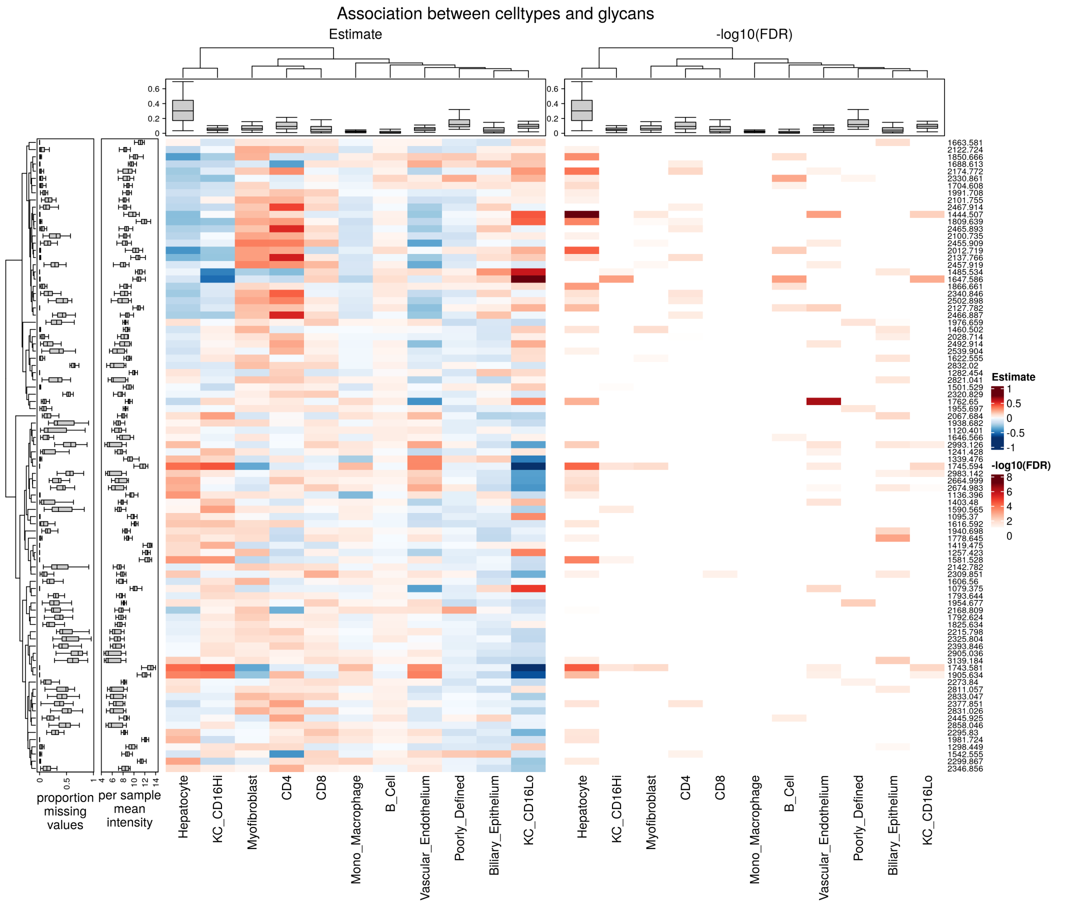


Figure S24: Results of statistical analysis, aggregated per sample. Heatmap per m/z and cell type for effect estimates (a) and -log10 FDR corrected p-values (b).

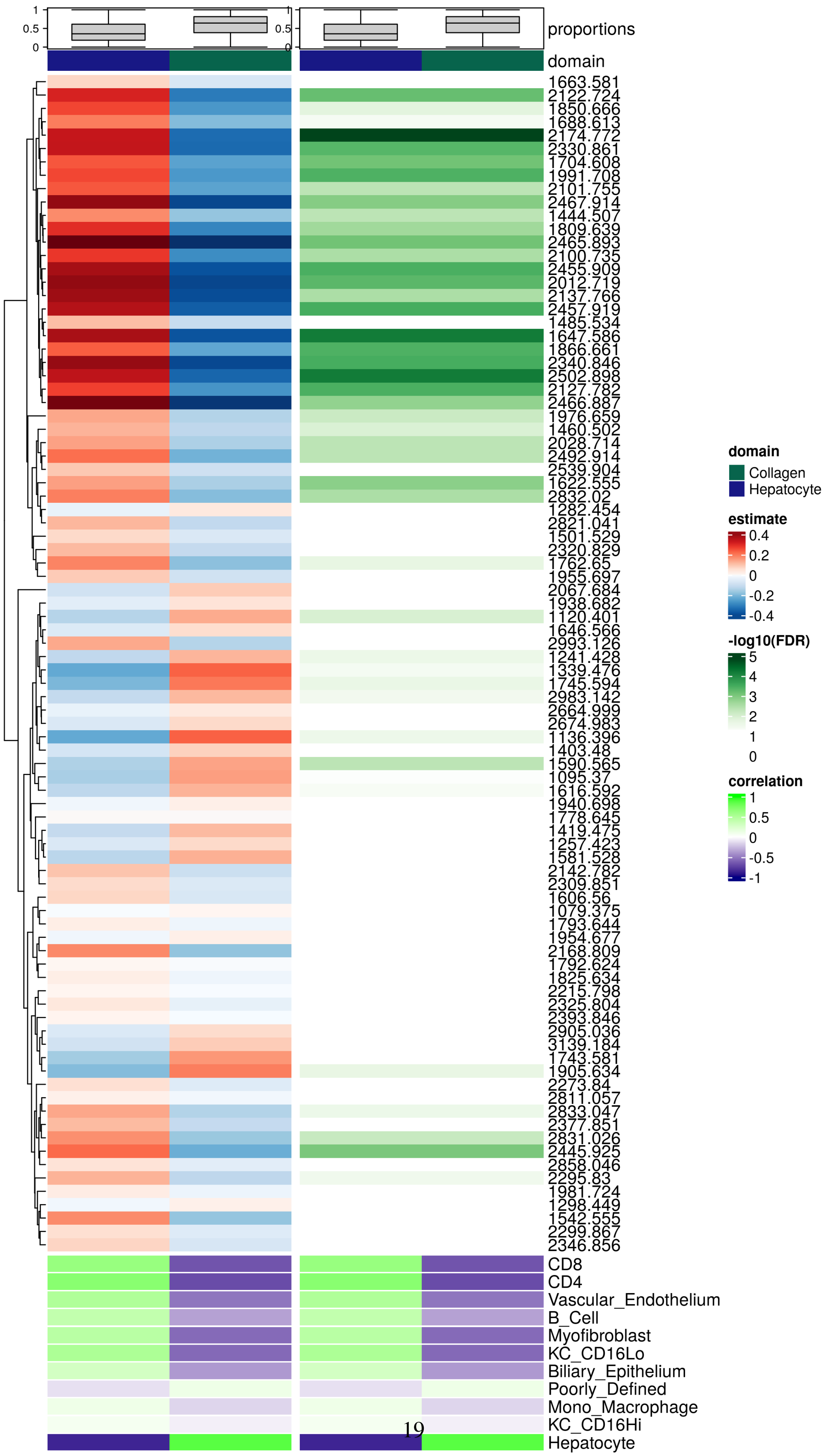


Figure S25: Results of statistical analysis, aggregated per domain and sample. Heatmap per m/z and domain for effect estimates (left) and -log10 FDR corrected p-values (right). Bottom Heatmap shows the correlation of cell type proportion with domain proportion.

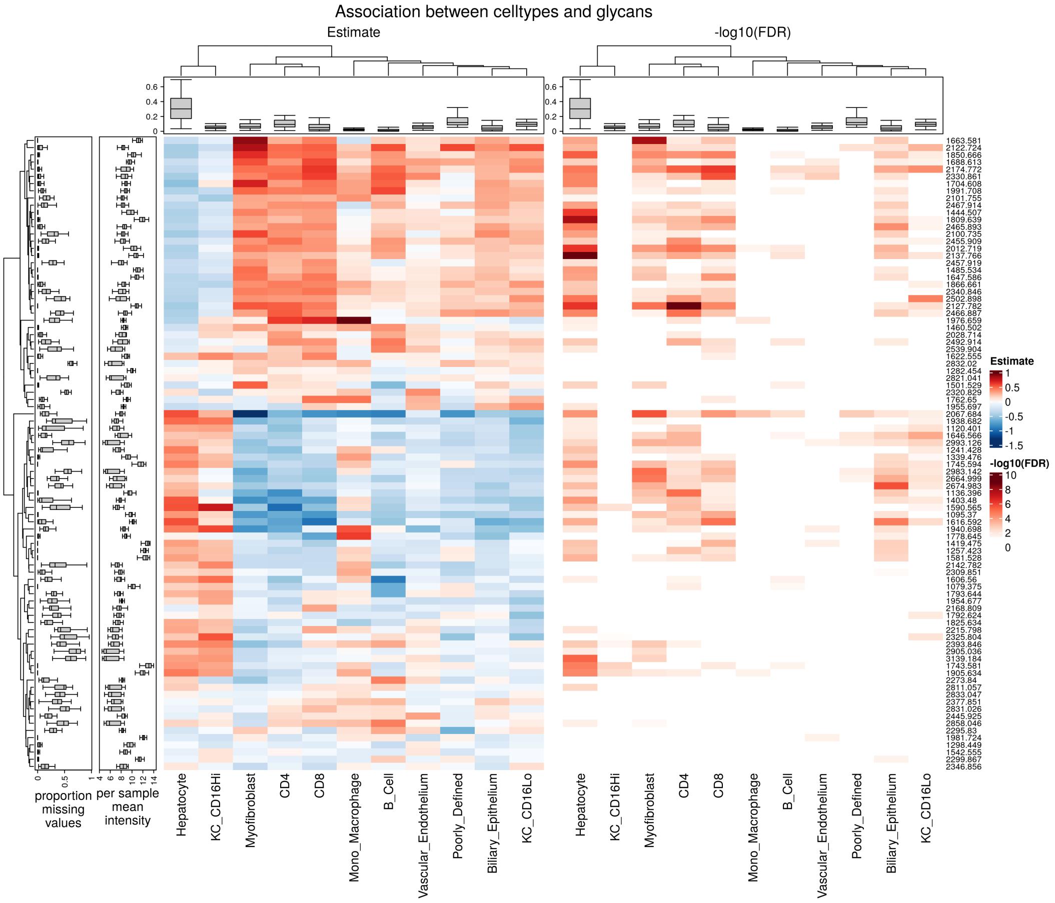


Figure S26: Results of statistical analysis. Heatmap per m/z and cell type for effect estimates (a) and -log10 FDR corrected p-values (b).

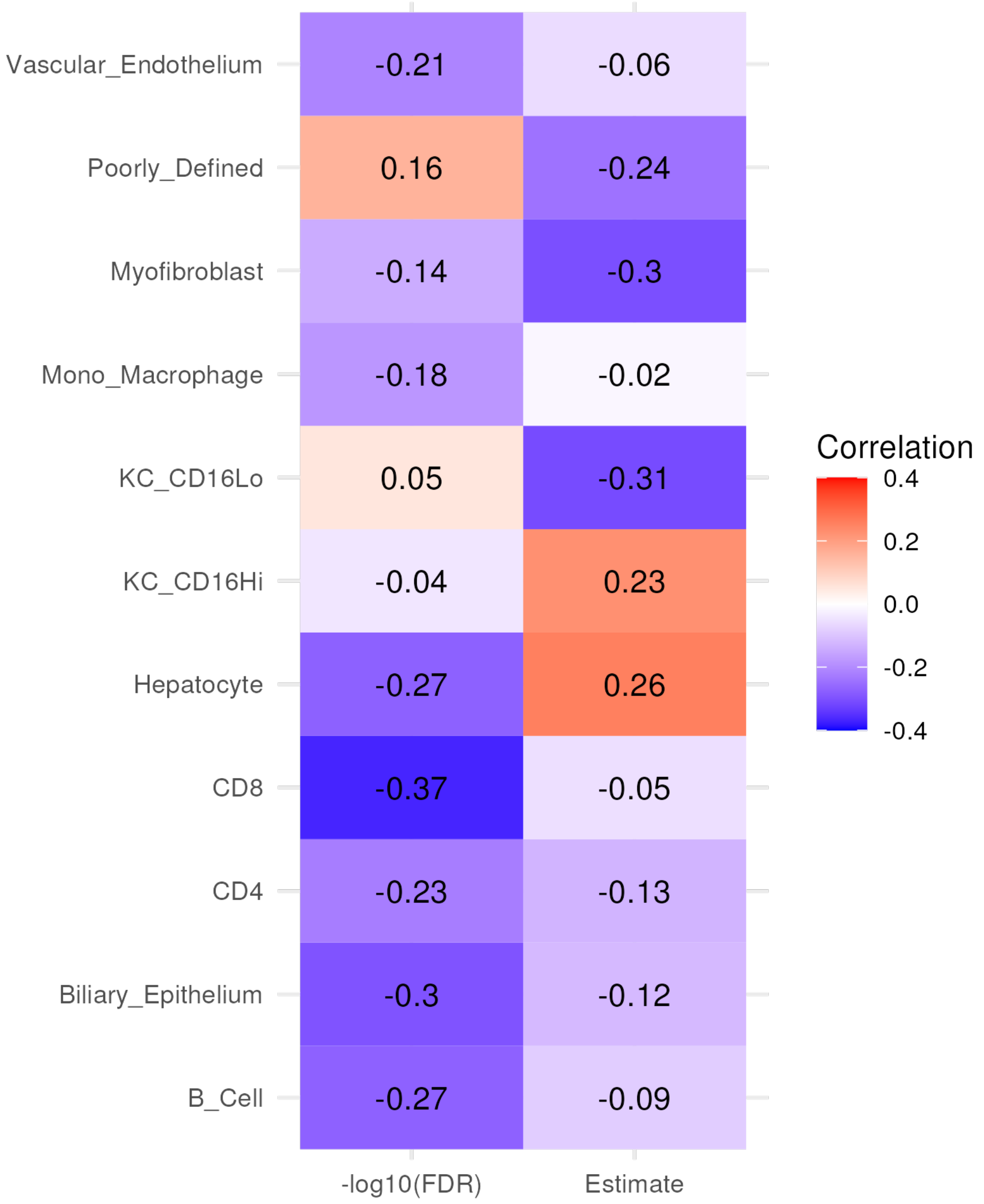


Figure S27: Heatmap of Correlations between either Estimates or p-values per cell type with proportion of missing m/z values.

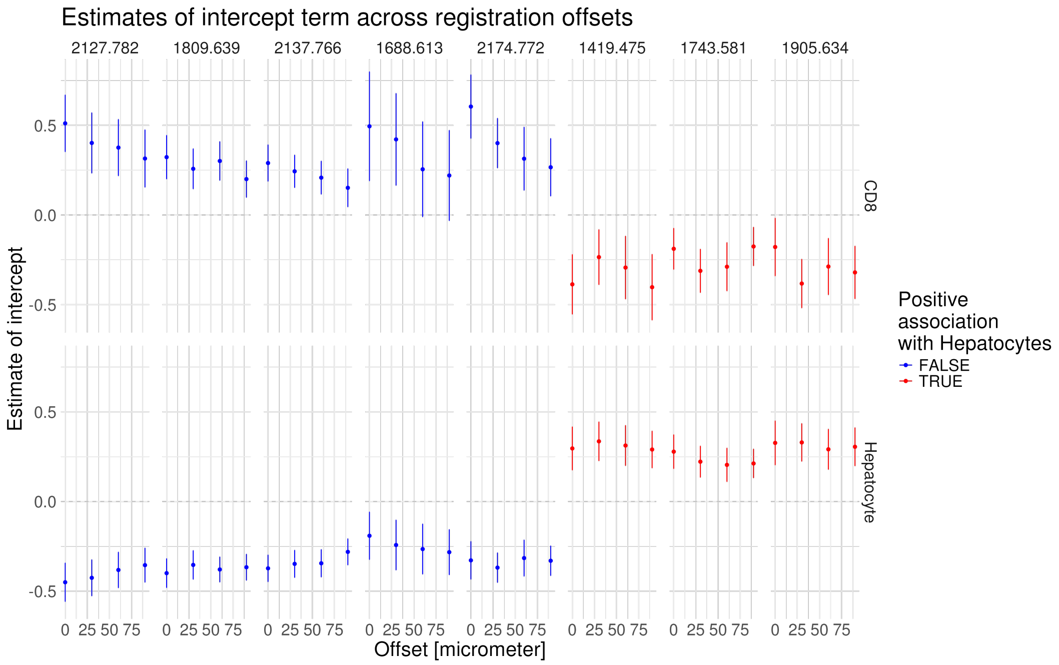


Figure S28: Sensitivity analysis of case study of m/z peaks with previously reported associations with either CD8 cells or Hepatocytes. Offset is the number of MSI pixels that are shifted in x-axis relative to the optimal registration. Shown are estimates and error bars ($+-1.96\times$ standard error).

| **Parameter** | **Value** |
|---|---|
| **Transfer** | |
| MALDI Plate Offset | 30.0 V |
| Deflection 1 Delta | 70.0 V |
| Funnel 1 RF | 450.0 Vpp |
| isCID Energy | 0.0 eV |
| Funnel 2 RF | 500.0 Vpp |
| Multipole RF | 500.0 Vpp |
| **Collision Cell** | |
| Collision Energy | 10.0 eV |
| Collision RF | 2900.0 Vpp |
| **Quadrupole** | |
| Ion Energy | 5.0 eV |
| Low Mass | m/z 700.00 |
| **Focus Pre TOF** | |
| Transfer Time | 110.0 $\mu s$ |
| Pre Pulse Storage | 10.0 $\mu s$ |

Table S3: Instrument Parameters

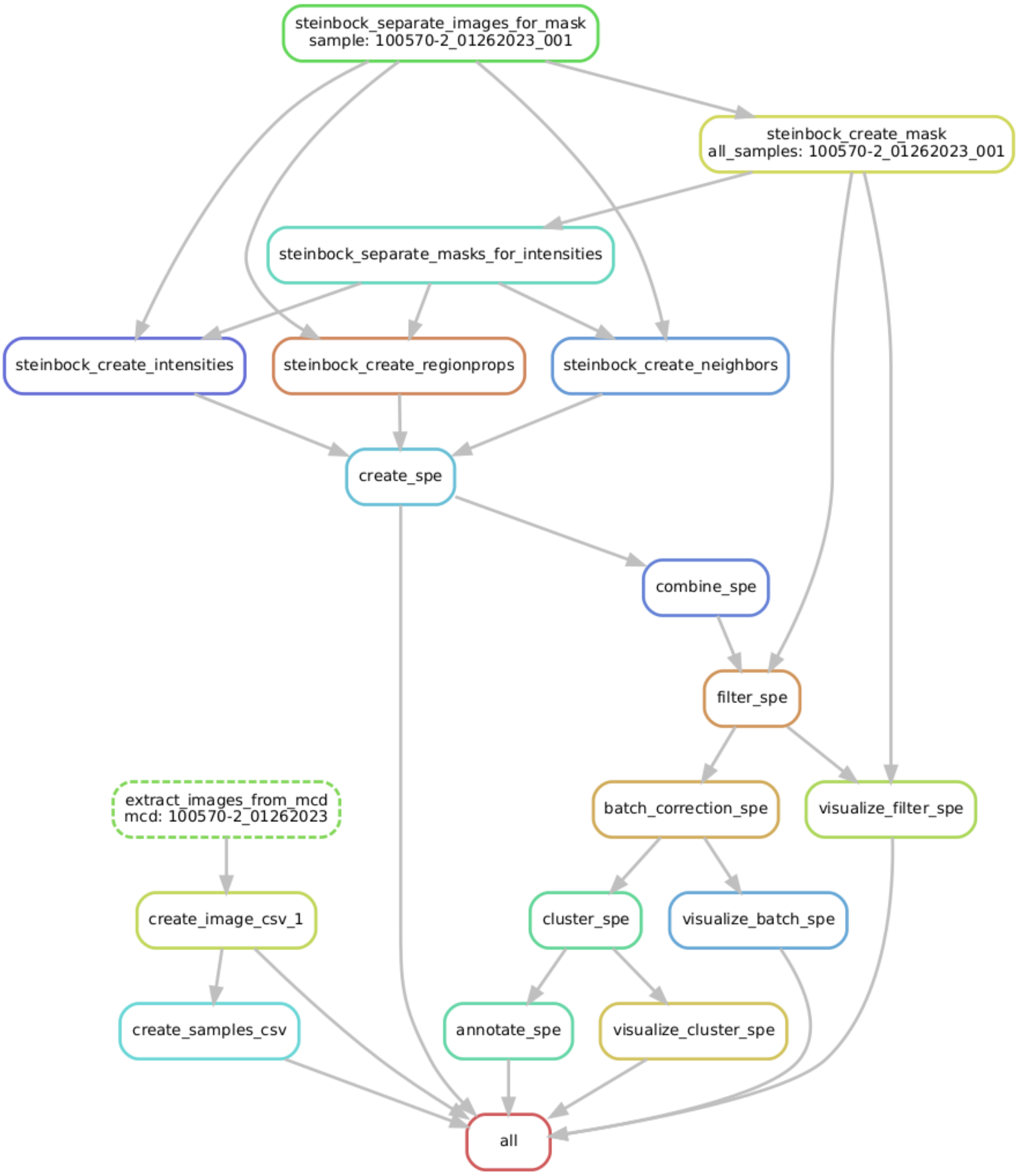


Figure S29: DAG of Snakemake workflow for IMC processing.

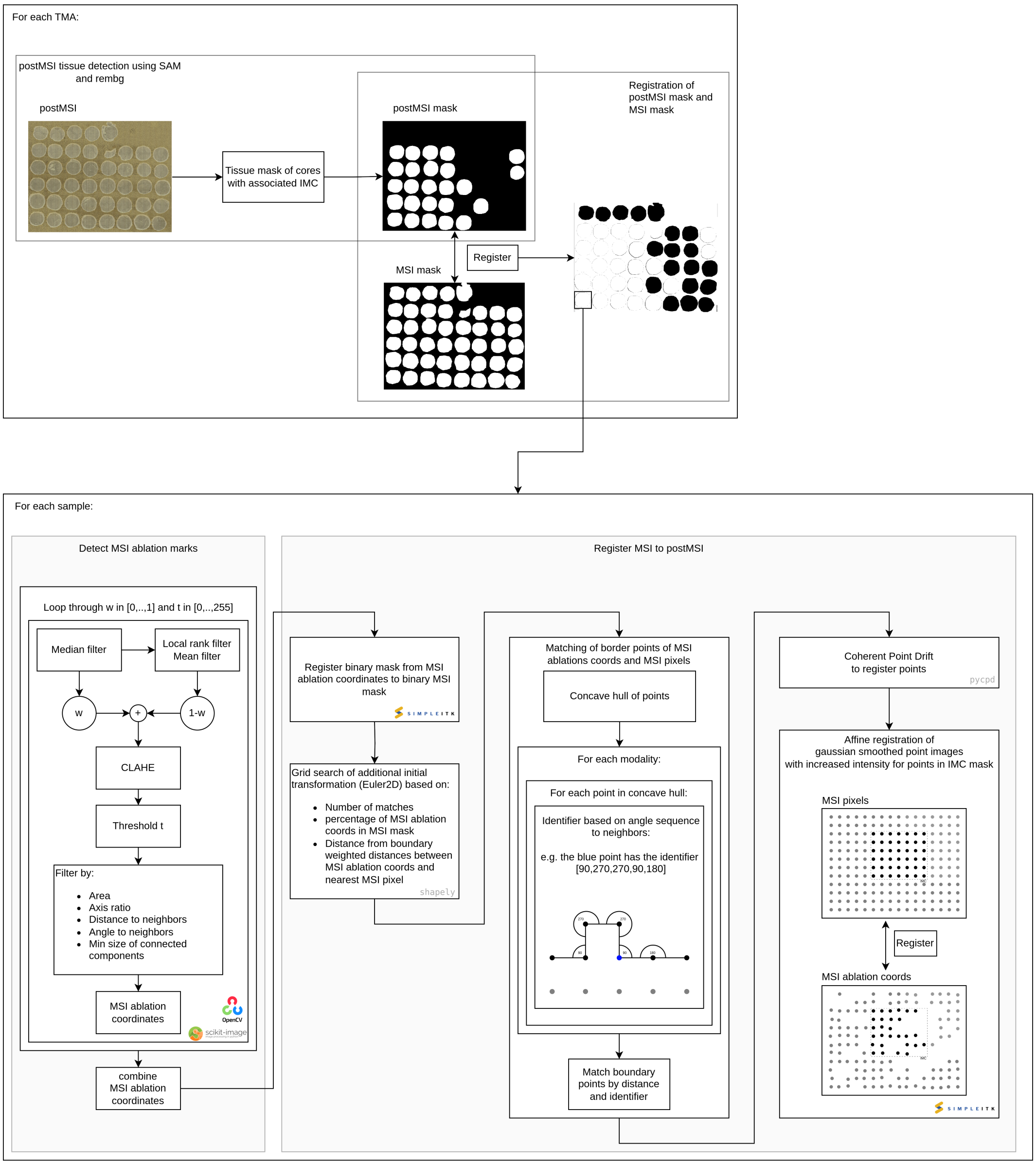


Figure S30: Diagram of the automatic registration of MSI to postMSI. First the TMA cores from the postMSI image and the MSI are matched (top panel). Next for each TMA core registration of MSI to postMSI ablation marks is done (bottom panels). After detection of the MSI ablation marks (bottom left panel) the detected marks are coregistered with the MSI pixels (bottom right panel).
